# Interactions of sequence diverse effector proteins of wheat powdery mildew control recognition specificity by the corresponding immune receptor

**DOI:** 10.1101/2024.12.30.629670

**Authors:** Jonatan Isaksson, Matthias Heuberger, Milena Amhof, Lukas Kunz, Salim Bourras, Beat Keller

## Abstract

To successfully colonize the living tissue of its host, the fungal wheat powdery mildew pathogen produces diverse effector proteins that are suggested to reprogram host defense responses and physiology. When recognized by host immune receptors, these proteins become avirulence (AVR) effectors. Several sequence-diverse AVRPM3 effectors and the suppressor of AVRPM3-PM3 recognition (SVRPM3^a1/f1^) are involved in triggering allele-specific, *Pm3*-mediated resistance, but the molecular mechanisms controlling their function in the host cell remain unknown. Here, we describe that AVRPM3^b2/c2^, AVRPM3^a2/f2^ and SVRPM3^a1/f1^ form homo- and heteromeric complexes with each other, suggesting they are present as dimers in the host cell. Alphafold2 modelling substantiated previous predictions that AVRPM3^b2/c2^, AVRPM3^a2/f2^ and SVRPM3^a1/f1^ all adopt a core RNase-like fold. We found that a single amino acid mutation in a predicted surface exposed region of AVRPM3^a2/f2^ resulted in recognition by the PM3b immune receptor, which does not recognize wildtype AVRPM3^a2/f2^. This indicates that differential AVRPM3 recognition by variants of the highly related PM3 immune receptors is due to subtle differences in similar protein surfaces of sequence-diverse AVRs. Based on our findings, we propose a model in which homodimers of AVRPM3s are recognized by their corresponding PM3 variants and that heterodimer formation with SVRPM3^a1/f1^ allows for evasion of recognition.

---

The wheat *Pm3* resistance gene comprises at least 17 functional alleles (*Pm3a–Pm3g*; *Pm3k– Pm3t*) that provide race-specific resistance to the powdery mildew fungal pathogen *Blumeria graminis* f.sp. *tritici* (*Bgt*) (Yahiaoui et al., 2004; Bhullar et al., 2009; Bourras et al., 2019). These alleles recognize effectors from *Bgt* to elicit effector-triggered immunity (ETI), resulting in hypersensitive response (HR)-mediated, localized cell death that limits pathogen growth (Balint-Kurti, 2019; Ngou et al., 2022). Despite over 97% amino acid sequence similarity, the *Pm3* alleles exhibit high specificity in recognizing their corresponding *AvrPm3* effectors (Bhullar et al., 2009, Bourras et al., 2015, 2019). In some cases, stronger and weaker pairs of *Pm3* alleles exist: the isolates recognized by the weak alleles, *Pm3f* and *Pm3c*, are completely encapsulated in the broader recognition spectrum of their corresponding stronger alleles *Pm3a* and *Pm3b*, respectively (Brunner et al., 2010).

Previous studies of the flax L5, L6 and L7 NLR-type immune receptors showed that they recognize variants of the same AVRL567 effector protein (Dodds et al., 2006). In contrast, the barley allelic variants MLA1, MLA6, MLA7, MLA10, MLA13 and MLA22 recognize the sequence diverse AVR_a1_, AVR_a6_, AVR_a7_, AVR_a10_, AVR_a13_ and AVR_a22_, respectively (Bauer et al., 2021; Lu et al., 2016; Saur et al., 2019). Structural studies revealed that AVR_a_ effectors share a common RNase-like fold which is conserved between different *Blumeria* species (Cao et al., 2023). The RNase-like fold consists of a core of several β-sheets facing a single α-helix and is hypothesized to have evolved from one common ancestor (Spanu, 2017). In *Bgt*, 70% or more of the 844 effectors are predicted to have structural similarity to the RNase-like fold (Cao et al., 2023; Müller et al., 2019). However, surprisingly little is known about how these highly similar effectors interact with each other.

Genetic analysis of avirulence in *Bgt* showed that multiple pathogen loci determine recognition by the *Pm3* alleles (Parlange et al., 2015, Bourras et al., 2015). The molecular identification of *AvrPm3* effectors revealed that AVRPM3^b2/c2^ is recognized by PM3b and PM3c, and AVRPM3^a2/f2^ is recognized by PM3a and PM3f (Bourras et al., 2015, 2019). Furthermore, a suppressor of avirulence (*Svr*) was identified as *SvrPm3^a1/f1^* which suppresses *Pm3* mediated recognitions of *AvrPm3* effectors (Bourras et al., 2015, 2019). The identification of these components showed that, like MLAs in barley, PM3 variants also recognize sequence diverse effectors with structural similarity in the form of an RNase-like fold (Bourras et al., 2019).

*SvrPm3^a1/f1^* acts as a central modifier of the *Pm3-AvrPm3* mediated resistance (Bourras et al., 2016). The level of suppression is dependent on the allele of *SvrPm3^a1/f1^* present in an isolate and correlated with its expression level (McNally et al., 2018). SVRPM3^a1/f1^ has the same core RNase-like fold commonly predicted for numerous *Bgt* and *Bh* effectors providing a possible explanation for its role as suppressor of plant immunity due to structural similarity with AVRPM3s. Here, we investigate the molecular interactions leading to the activation or the suppression of the AVRPM3-PM3 effector-NLR pairs. In particular, we explore the possible role of effector-effector interactions and structural variation as mechanisms controlling specificity of AVRPM3 recognition and suppression of PM3-mediated resistance.

To investigate whether multimer formation plays a role in the functionality of the AVRPM3 effectors, we first tested AVRPM3^b2/c2^ for interactions with itself. The signal peptide of AVRPM3^b2/c2^-FLAG was removed and the resulting protein was transiently expressed in *Nicotiana benthamiana* through Agrobacterium-mediated infiltration. Immunoblot analysis of whole cell lysates applied to sodium dodecyl-sulfate polyacrylamide gel electrophoresis (SDS-PAGE) under denaturing conditions revealed bands at multiple molecular weights: a 16 kilodalton (kDa) band corresponding to the predicted size of the AVRPM3^b2/c2^ monomer, as well as 32 kDa and 48 kDa bands at the predicted size of a dimer and trimer, respectively (Fig. 1a). These observations suggest that AVRPM3^b2/c2^ may form higher-order complexes *in planta*.

**Figure 1.**
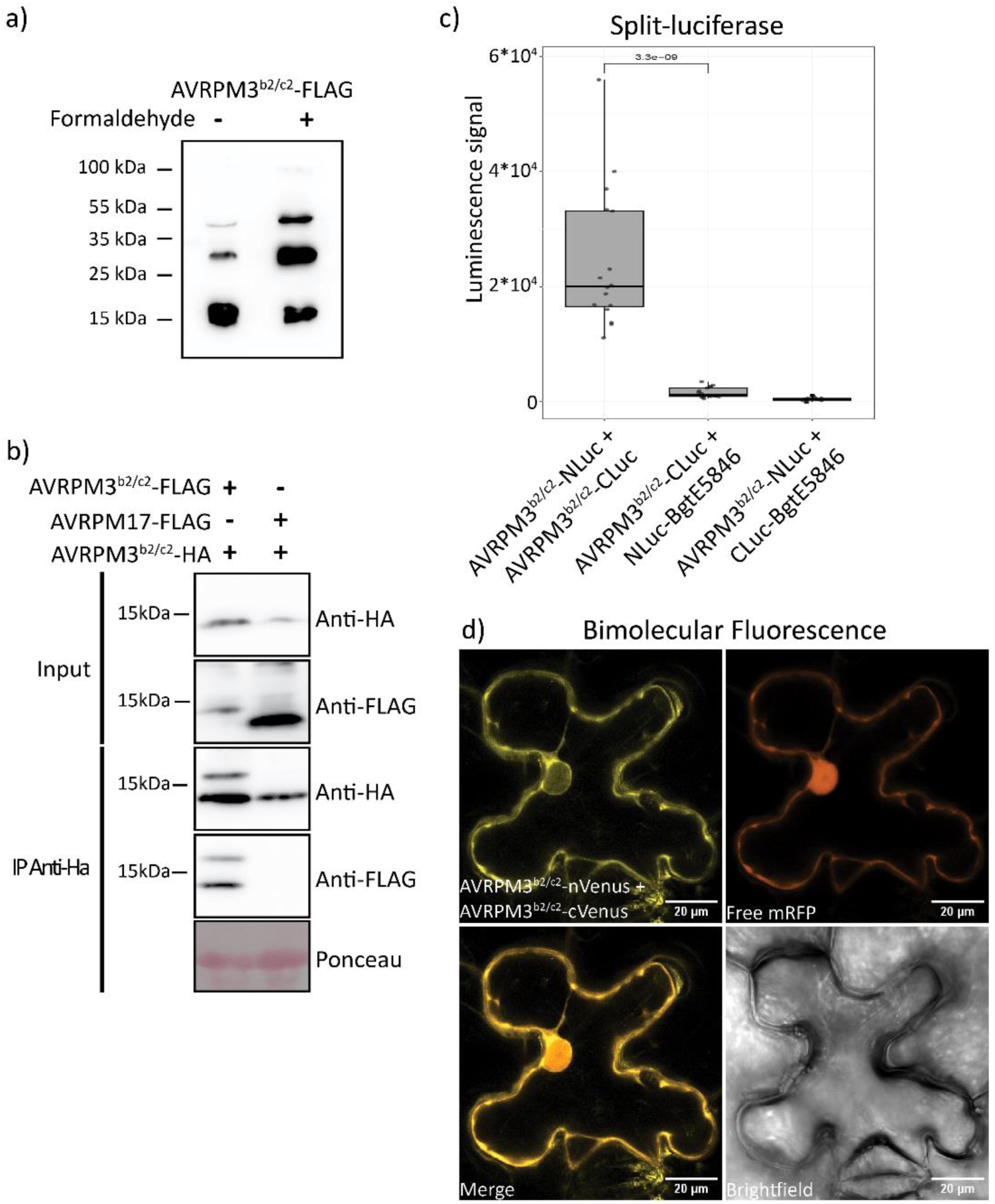
Homodimer formation of AVRPM3^b2/c2^. (a) Immunoblot of AVRPM3^b2/c2^-FLAG protein lacking the signal peptide and transiently expressed via Agrobacterium-mediated transformation in *Nicotiana benthamiana*. AVRPM3^b2/c2^-FLAG was detected at the predicted size of 16kDa and higher bands with the size of ∼32kDa and ∼48kDa. Formaldehyde crosslinking of the tissue is indicated by a + sign and shows an increase in the bands at the larger sizes of ∼32kDa and ∼48kDa (b) Immunoblot of co-immunoprecipitation experiment of effector proteins lacking the signal peptide and transiently expressed via Agrobacterium-mediated transformation in *N. benthamiana*. Co-expression of AVRPM3^b2/c2^-HA with AVRPM3^b2/c2^-FLAG or AVRPM17-FLAG followed by anti-HA magnetic bead pulldown. Antibodies used for immunoprecipitation (IP) and immunoblotting as indicated. (c) Split-luciferase assay between AVRPM3^b2/c2^ and itself in *N. benthamiana*. AVRPM3^b2/c2^-Cluc with Nluc-BgtE-5846 and AVRPM3^b2/c2^-Nluc with Cluc-BgtE-5846 served as controls for specific interaction between AVRPM3^b2/c2^ and itself. Luminescence signal was measured three days post infiltration (dpi). In the plot, data points refer to four independent biological replicates, each with four technical replicates. Statistical differences between the test and control were assessed using an un-paired Wilcoxon rank sum test with the p-value indicated. (d) Bimolecular Fluorescence Complementation Assay (BiFC) shows the interaction between AVRPM3^b2/c2^ and itself while co-expressed with a free mRFP in *N. benthamiana*. Fluorescence signal was detected via laser scanning confocal microscopy at 3 dpi. The scale bar (in white) for all images is 20μm.

To further validate dimers and multimers, tissue cross-linking was performed using formaldehyde before whole cell lysate preparation (see Methods). Indeed, upon cross-linking the AVRPM3^b2/c2^ band from the monomer shifted towards the dimeric and trimeric forms (Fig. 1a). We therefore conclude that AVRPM3^b2/c2^ is prone to multimerization which is stabilized through cross-linking and robust enough to be maintained under denaturing conditions.

To investigate if AVRPM3^b2/c2^ forms homodimers specifically *in planta,* additional approaches were used, including co-immunoprecipitation (co-IP), split-luciferase (Gehl et al., 2011), and bimolecular fluorescence complementation (BiFC) assays. In the co-immunoprecipitation experiment, C-terminally FLAG- and HA-tagged versions of AVRPM3^b2/c2^ were co-expressed in *N. benthamiana*. Immunoprecipitation with anti-HA magnetic beads revealed that AVRPM3^b2/c2^-HA co-immunoprecipitated AVRPM3^b2/c2^-FLAG, indicating homodimer formation within the plant (Fig. 1b). As a negative control, we used AVRPM17, a *Bgt* effector predicted to belong to the same RNase-like fold family. It is specifically recognized by PM17, a homolog of PM3 (Singh et al., 2018). Consistent with specific AVRPM3^b2/c2^ homodimer formation, co-immunoprecipitation assays with AVRPM3^b2/c2^-HA revealed no detectable levels of AVRPM17-FLAG in the immunoblot (Fig. 1b).

We further validated AVRPM3^b2/c2^ homodimerization through a split-luciferase assay, where full-length AVRPM3^b2/c2^ constructs were fused with either the N-terminal (NLuc) or C-terminal (CLuc) domains of luciferase and co-expressed in *N. benthamiana*. As a control, we used BgtE5846, an RNase-like effector which was previously used in FLuCI experiments (Manser et al., 2024). Consistent with the co-IP assays, strong luminescence was observed when AVRPM3^b2/c2^-NLuc and AVRPM3^b2/c2^-CLuc were co-expressed, indicative of AVRPM3^b2/c2^ dimerization (Fig. 1c). In contrast, significantly lower luminescence was observed in control interactions using BgtE5846. These results further supported the specificity of AVRPM3^b2/c2^ dimerization.

Lastly, the dimerization of AVRPM3^b2/c2^ was also tested using BiFC. There, AVRPM3^b2/c2^ was fused with either the N-terminal (nVenus) or C-terminal (cVenus) half of the Venus fluorescent protein. Co-expression of nVenus- and cVenus-tagged AVRPM3^b2/c2^ in *N. benthamiana* produced a fluorescence signal, further confirming the formation of a homodimer (Fig. 1d). Free mRFP was also co-expressed with the BiFC constructs as a marker to distinguish the cytoplasmic and nuclear compartments of the cell. The fluorescence signal indicated that the AVRPM3^b2/c2^ dimer was primarily localized to the cytoplasm with faint nuclear localization as indicated by the restricted co-localization with the free mRFP marker (Fig. 1d). Together these experimental observations indicate that AVRPM3^b2/c2^ forms a dimer *in planta* and the limited nuclear signal suggests that AVRPM3^b2/c2^ is primarily cytoplasmic.

Interestingly, Qi and colleagues (2024) also found that effector candidates of the biotrophic pathogen *Phakospora pachyrhizi* form a complex network of interactions with each other, suggesting that effector multimerization occurs frequently. While interactions between AVRPM3 effectors and other *Bgt* effectors cannot be excluded, AVRPM3^b2/c2^ did not interact with AVRPM17 or BgtE5846. Therefore, despite sharing a predicted structure and ancestry with these effectors (Bourras et al., 2019; Manser et al., 2021; Müller et al., 2022; Spanu, 2017), AVRPM3^b2/c2^ appears to have diverged sufficiently from them to not interact.

We then examined SVRPM3^a1/f1^ for its ability to form homodimers or interact with AVRPM3^b2/c2^ and AVRPM3^a2/f2^. Our reasoning was that interactions between SVRPM3^a1/f1^ and the AVRPM3 effector could provide new insights on the mode of suppression of the AVRPM3-PM3 interaction. We used SVRPM3^a1/f1^ variant from Bgt_94202, a mildew isolate which has been shown to be virulent towards all *Pm3* alleles. We also used a variant encoded by Bgt_96224, an isolate shown to be avirulent towards all *Pm3* alleles (Bourras et al., 2015). All constructs were produced with either FLAG or HA tags and expressed in *N. benthamiana* to detect possible SVRPM3^a1/f1^ homodimers, or heterodimers between SVRPM3^a1/f1^ variants and AVRPM3^b2/c2^ or AVRPM3^a2/f2^.

Following co-immunoprecipitation with anti-FLAG magnetic beads, we found that both SVRPM3^a1/f1^ variants co-immunoprecipitated with themselves, AVRPM3^b2/c2^ and AVRPM3^a2/f2^, indicating both homo and heterodimer formation (Fig. 2a). Furthermore, neither variant of SVRPM3^a1/f1^ interacted with AVRPM17 indicating that the interaction between SVRPM3^a1/f1^ and the AVRPM3 effectors is specific (Fig. 2a). Interestingly, SVRPM3^a1/f1^ variants were more enriched if compared to AVRPM3 effectors in our co-immunoprecipitation experiment (Fig. 2a). This suggests that SVRPM3^a1/f1^ has a strong affinity to interact with itself and this self-association leads to a reduced association of SVRPM3^a1/f1^ with AVRPM3s. However, further testing to determine affinities between the effectors is needed to quantitatively assess whether this is the case.

**Figure 2.**
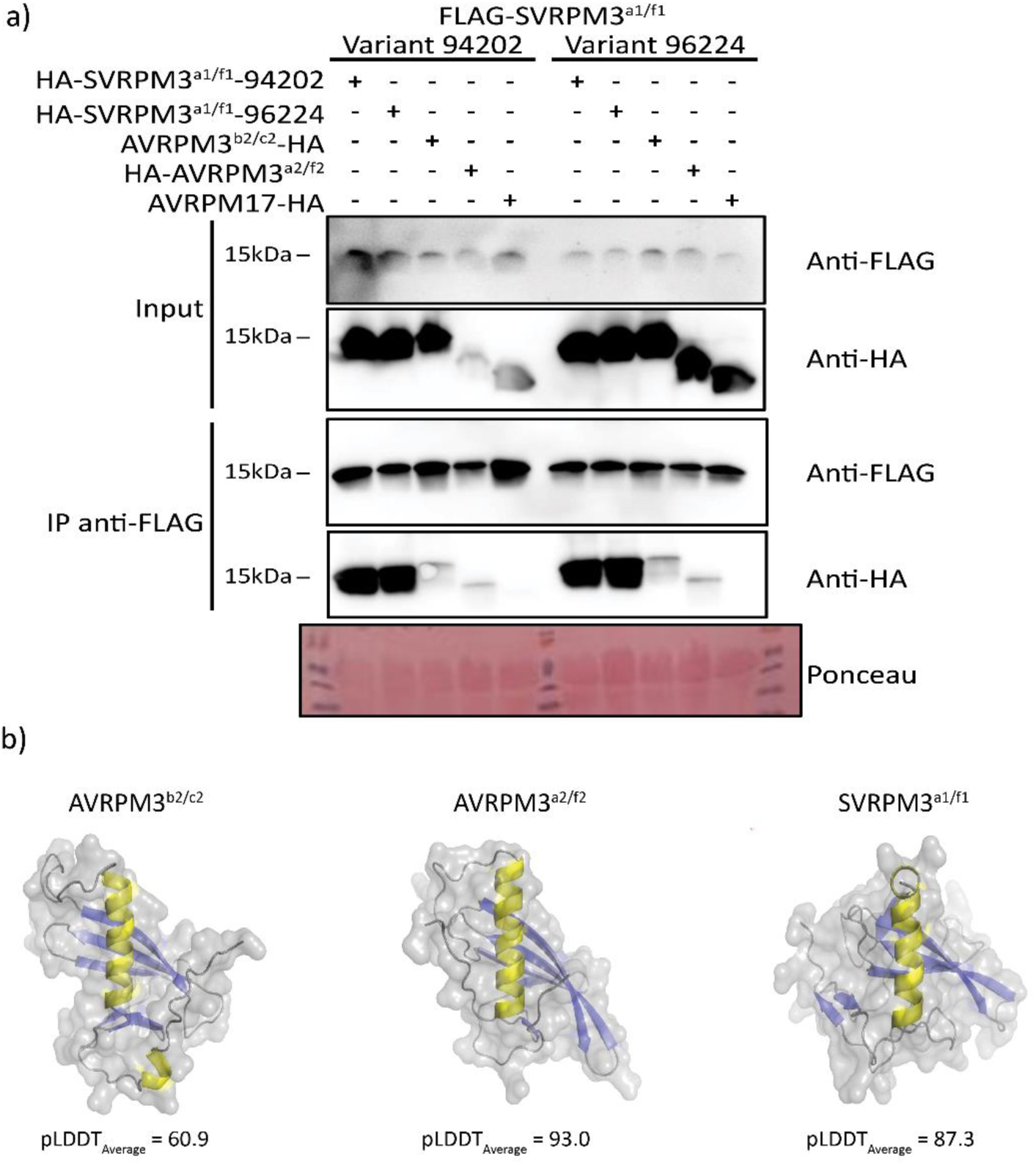
SVRPM3^a1/f1^ homodimerizes and forms heterodimer complexes with two different AVRPM3 effectors with which it shares a conserved core structure. (a) Immunoblot of co-immunoprecipitation experiment of effector proteins lacking the signal peptide and transiently expressed via Agrobacterium-mediated transformation in *Nicotiana benthamiana*. FLAG-SVRPM3^a1/f1^ variants from *Bgt* isolates 94202 and 96224 co-expressed with HA-SVRPM3^a1/f1^-94202, HA-SVRPM3^a1/f1^-96224, AVRPM3^b2/c2^-HA, HA-AVRPM3^a2/f2^ or AVRPM17-HA. Tissue was harvested 2-3 days post infiltration and followed by a FLAG magnetic bead pulldown. Antibodies used for immunoprecipitation (IP) and immunoblotting as indicated. (b) cartoon and surface representation of the top ranked Alphafold2 models of AVRPM3^b2/c2^, AVRPM3^a2/f2^ and SVRPM3^a1/f1^ with their pLDDT scores as indicated. Residues colored in yellow correspond to α-helices while those in blue correspond to β-sheets.

Previous studies showed that *AvrPm3^a2/f2^*, *AvrPm3^b2/c2^*, and *SvrPm3^a1/f1^* are present in all tested *Bgt* isolates globally despite the selection pressure to escape detection by PM3 variants (Bourras et al., 2019; McNally et al., 2018). This suggests that cooperation between these effectors might be beneficial for growth, potentially through hetero- and homodimer complexes related to their virulence functions. Specific interaction between SVRPM3^a1/f1^ and AVRPM3s might have evolved because they have converged on similar functions or targets in the host, which would separate them from the other effectors tested in this study such as AVRPM17 or BgtE5846 which they did not interact with despite also being predicted to have an RNase-like fold. This aligns with recent experimental and computational observations that inner core residues in fungal effectors such as the RNase-like fold are conserved, while surface-exposed residues diversify to drive functional adaptation (Cao et al., 2024; Derbyshire and Raffaele, 2023).

To validate previous findings and provide an improved structural model for SVRPM3^a1/f1^, AVRPM3^a2/f2^ and AVRPM3^b2/c2^ we applied all proteins to structure prediction using Alphafold2 and Alphafold3 (Jumper et al., 2021). We found that predictions with Alphafold2 resulted in more robust structures, that were better supported than those obtained by Alphafold3 based on the pLDDT scores (Supplementary table 1). All three effectors displayed a conserved core RNase-like fold, with a α-helix surrounded by β-sheets (Fig. 2b). All models generated a predicted local distance difference test (pLDDT) score >50, indicating that they were robust (Jumper et al., 2021). The AVRPM3^a2/f2^ and SVRPM3^a1/f1^ predicted structures had scores of 93.0 and 87.3, respectively. The score for AVRPM3^b2/c2^ was 60.9 which is near the threshold and therefore represents a poorer model than AVRPM3^a2/f2^ and SVRPM3^a1/f1^ models (Fig. 2b). These results suggest that SVRPM3^a1/f1^, AVRPM3^a2/f2^ and AVRPM3^b2/c2^ share a RNase-like fold, and we propose that such structural relatedness may explain the capacity of these effectors to form heterodimers.

Considering the structural similarity between AVRPM3^b2/c2^ and AVRPM3^a2/f2^ we decided to experiment with the hypothesis that small changes in the predicted surface exposed amino acids of the AVRPM3 effectors could lead to cross-recognition by a non-corresponding PM3 variant. To test this, we used synthetic effector protein variants encoding single amino acid polymorphisms as compared to wildtype AVRPM3^a2/f2^ (a.k.a. variant A) which were previously described by McNally and collaborators (2018). The single amino acid changes in AVRPM3^a2/f2^ were based on the variation found within its effector family with the logic that conserved residues would be important for their structure and therefore lead to conservation of important structural elements (McNally et al., 2018) such as those found within their core RNase-like fold (Fig. 2b).

We focused on eight variants which were shown to either (i) induce stronger HR response as compared to wildtype AVRPM3^a2/f2^ when co-expressed with PM3a or (ii) harbored mutations in a larger region suggested to specifically act on the strength of the HR response induced by PM3A upon AVRPM3^a2/f2^ recognition (McNally et al., 2018). The mutations in the AVRPM3^a2/f2^ variants that were tested against PM3b can be found in the schematic depiction of AVRPM3^a2/f2^ (Fig. 3a). The variants were re-tested against PM3a and showed a similar pattern to that observed previously (Supplementary fig. 1a; McNally et al., 2018). With the exception of AVRPM3^a2/f2^-L90E, the other seven AVRPM3^a2/f2^ variants triggered HR when combined with PM3a, with the strongest HR seen for AVRPM3^a2/f2^-L91Y (Supplementary fig. 1a).

**Figure 3.**
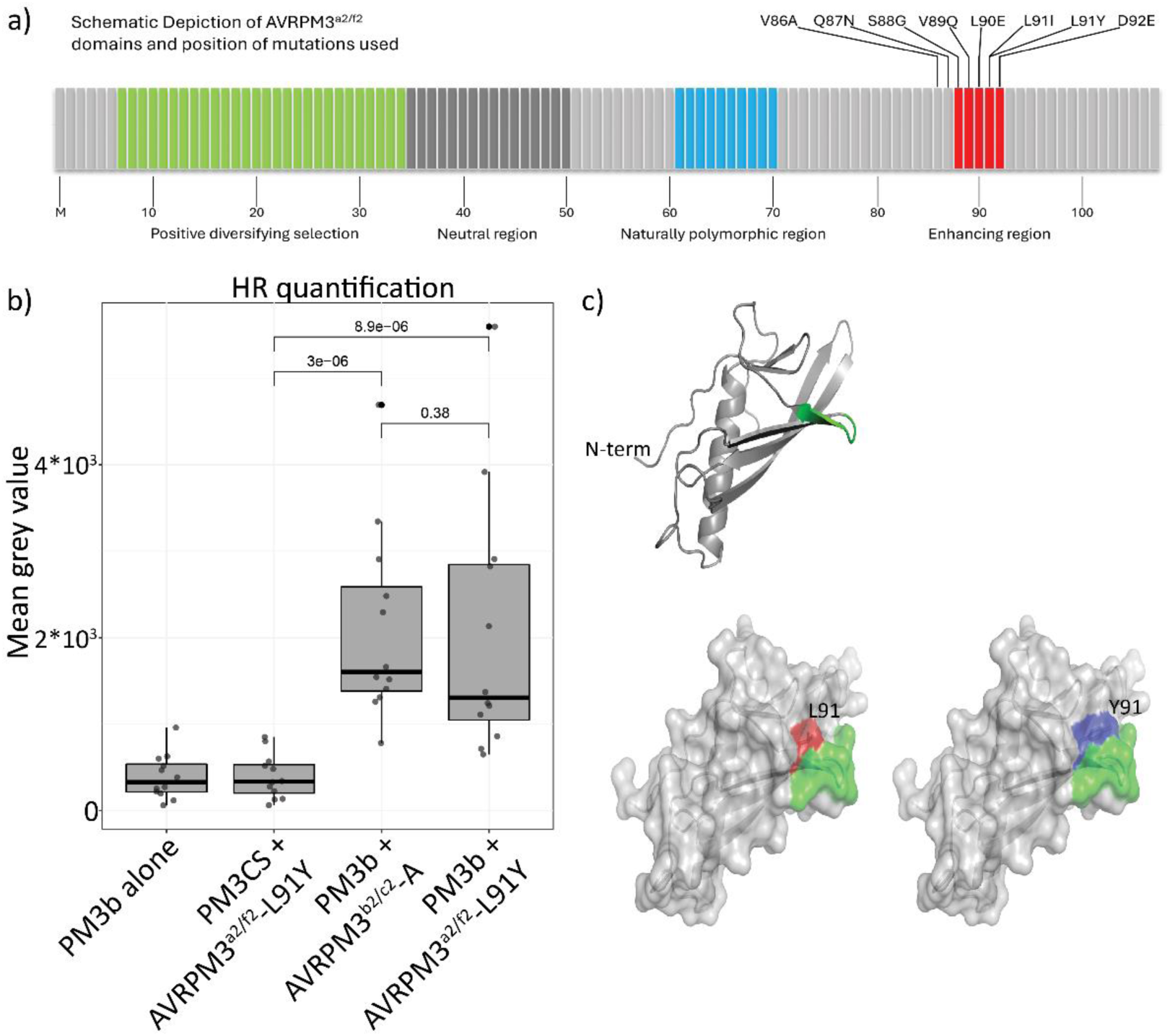
A single L91Y amino acid substitution in AVRPM3^a2/f2^-A leads to recognition by the non-corresponding PM3b. (a) schematic depiction of AVRPM3^a2/f2^ showing the sites of the single residue mutations as compared to variant A used in the cross-recognition screen. Sites previously identified to be under diversifying selection, neutral, naturally polymorphic and enhancing of HR are depicted below. Adapted from McNally et al., 2018. (b) Transient Agrobacterium mediated transformation of *N. benthamiana* leaves of PM3b with variant A of AVRPM3^b2/c2^ or with AVRPM3^a2/f2^-L91Y lacking their signal peptides. PM3b alone and PM3CS (a susceptible variant) co-expressed with AVRPM3^a2/f2^-L91Y were used as negative controls. Fluorescence signal due to cell death was measured 5 days post infiltration with n=12 biological replicates. P-values were calculated using an un-paired Wilcoxon rank sum test. (c) Cartoon and surface representation of the top ranked models from Alphafold2 of AVRPM3^a2/f2^-A and AVRPM3^a2/f2^-L91Y (pLDDTAverage = 93.3, pLDDT of L/Y = 91/94). The residues of the putative enhancing/recognition region of AVRPM3^a2/f2^-A are colored in green in all three models (cartoon and surfaces). The L91 residue in AVRPM3^a2/f2^-A is surface colored in red while the Y91 residue of AVRPM3^a2/f2^-L91Y is surface colored in blue.

Interestingly, our screen for cross-recognition revealed that AVRPM3^a2/f2^-L91Y demonstrated a unique ability to trigger HR also with PM3b, which does not recognize wildtype AVRPM3^a2/f2^ (Supplementary fig. 1b). Furthermore, the HR triggered by AVRPM3^a2/f2^-L91Y was as strong as the one triggered by AVRPM3^b2/c2^, i.e. the corresponding AVR of PM3b (Fig. 3b). Lastly, the HR triggered by PM3b was specific, as neither the susceptible variant PM3CS nor the PM3d and PM3e variants triggered HR with AVRPM3^a2/f2^-L91Y (Fig. 3b; Supplementary fig. 2a). Structural modeling of AVRPM3^a2/f2^-L91Y showed that the substitution of leucine with tyrosine at position 91 introduced a surface-exposed protrusion in the protein structure (Fig. 3c). We thus conclude that AVRPM3^a2/f2^-L91Y-PM3b is a genuine case of AVR cross-recognition by a non-corresponding NLR through a single residue change in AVRPM3^a2/f2^.

Recently, it was shown that the inactive form of PM3b interacts most strongly with AVRPM3^b2/c2^, suggesting that activation weakens the effector-receptor complex (Isaksson et al., 2024). Whether dimerization of AVRPM3 effectors is essential for recognition by PM3 variants offers an exciting avenue for further exploration. Based on our findings, we propose a model in which AVRPM3^b2/c2^ and AVRPM3^a2/f2^ are recognized as homodimers by their corresponding PM3 variants (Fig. 4), Furthermore, this is likely to occur in a similar manner based on the finding that a single amino acid polymorphism can extend AVRPM3^a2/f2^ recognition to the non-corresponding PM3b (Fig. 3b). Based on the observed heterodimer formation of the AVRPM3s with SVRPM3^a1/f1^, we hypothesize that this alters effector recognition and enables evasion of *Pm3*-mediated resistance. In a previous study, it was shown that the extent to which SVRPM3^a1/f1^ can suppress PM3 mediated resistance is strongly associated with the variant present and its expression level (Bourras et al., 2015). Interestingly, we found that both versions of SVRPM3^a1/f1^ could interact with AVRPM3^b2/c2^ and AVRPM3^a2/f2^ (Fig. 2a) despite SVRPM3^a1/f1^-96224 coming from an avirulent isolate (Bourras et al., 2015). Therefore, according to our proposed model, SVRPM3^a1/f1^-96224’s inability to suppress AVRPM3 recognition by PM3 in wheat might be due to a combination of lowered activity due to differences imposed by the variant amino acids, as well as its lower expression levels as compared to *SvrPm3^a1/f1^-94202*.

**Figure 4.**
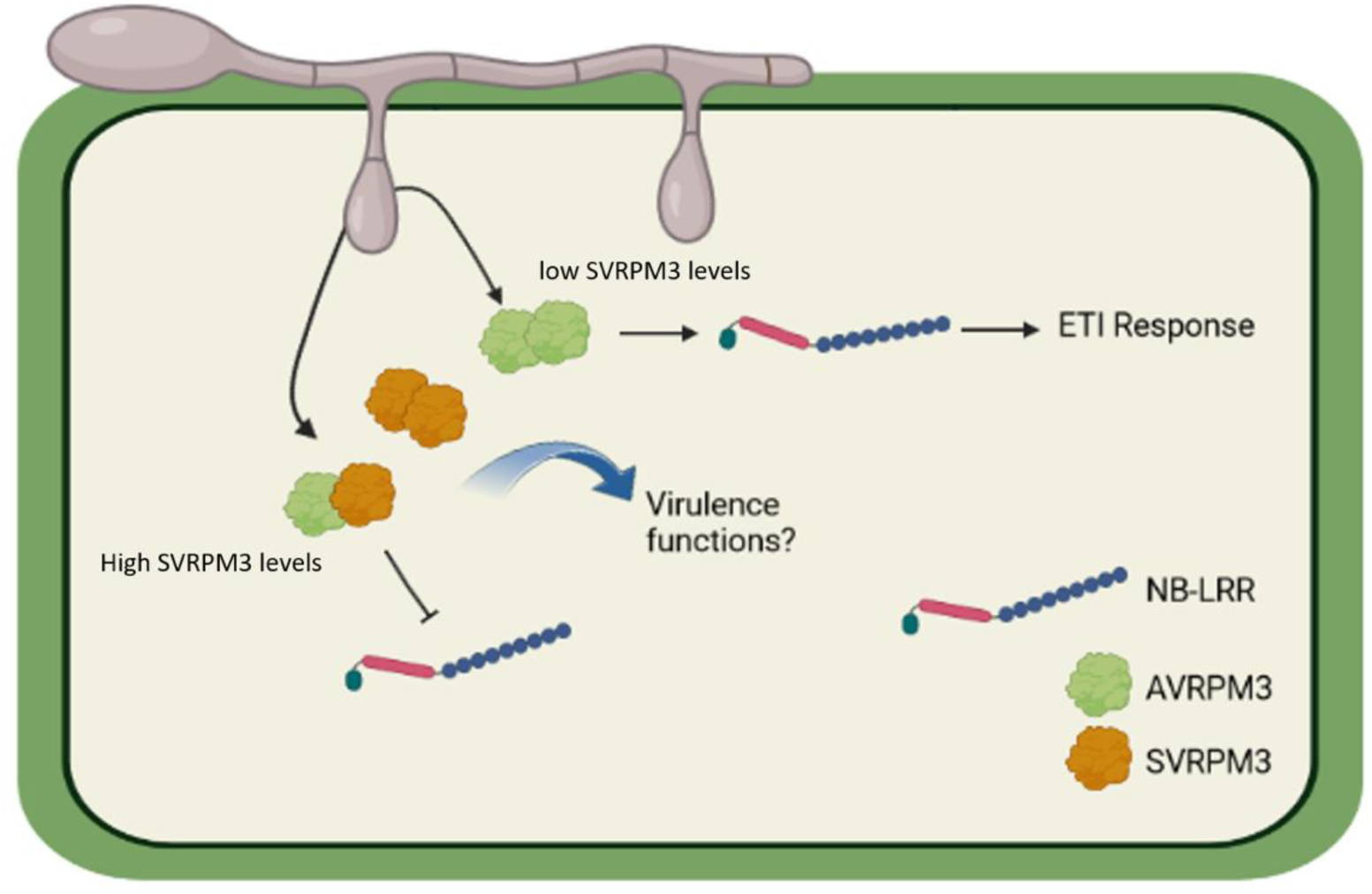
Model of AVRPM3 and SVRPM3^a1/f1^ effector interactions determining the outcome of avirulence. AVRPM3 effectors and SVRPM3^a1/f1^ are secreted from the fungus into the cytoplasmic space. When the ratio of AVRPM3 effector to SVRPM3^a1/f1^ is high, the concentration of the suppressor is insufficient to outcompete the homodimer formation, thus leading to recognition of the correctly formed homodimer of AVRPM3 by its corresponding PM3 variant which initiates an ETI response. If there is a sufficient amount of SVRPM3^a1/f1^, the heteromeric complex is formed and there is no activation of PM3-mediated signaling. The homodimer and heterodimer complexes might all have a function in virulence. Model created using Biorender.

In summary, our goal was to better understand the molecular interaction of the fungal effectors involved in the AVRPM3–PM3–SVRPM3 interaction and their contributions to allele-specific resistance by *Pm3*. Our results underline the significance of structural conservation over sequence similarity of AVRPM3 effectors in PM3 mediated resistance. This study provides a possible mechanism for how fungal effectors achieve virulence and contribute to resistance evasion, a crucial consideration for developing strategies to enhance plant immunity.

## Materials and methods

Materials and methods used in this study can be found in supplementary note 1.

## Acknowledgments

Imaging was performed with equipment maintained by the Center for Microscopy and Image Analysis, University of Zurich. This work was supported by grants 310030_204165 and 310030B_182833 from the Swiss National Science Foundation.

## Author contributions

J.I., S.B., and B.K. designed the research. J.I., M.H., L.K. and M.A. performed the experiments. J.I. analyzed the data. J.I., L.K., S.B., and B.K. wrote the manuscript.

## Supplementary materials sample

**Supplementary note 1**

**Supplementary table 1**

**Supplementary figure 1**

**Supplementary figure 2**

**References**

