## Supplementary Material for "Interactions of sequence diverse effector proteins of wheat powdery mildew control recognition specificity by the corresponding immune receptor"

Contents:

**Supplementary note 1**

**Supplementary table 1**

**Supplementary figure 1**

**Supplementary figure 2**

**References**

**Supplementary note 1**

**Materials and Methods**

**Transient expression in *Nicotiana benthamiana***

*Agrobacterium tumefaciens* transformed with binary vectors containing the construct of interest were prepared in Luria broth (LB) containing the appropriate antibiotics and incubated overnight at 28°C with 200 rpm shaking. Overnight cultures were harvested by centrifugation at 3,300xg for 5 minutes and washed by re-suspending in fresh LB media without antibiotics. Cells were harvested again by centrifugation at 3,300xg for 5 minutes and re-suspended in a small amount of AS-medium (10mM MES-KOH, pH5.6; 10mM $MgCl_2$; 100µM acetosyringone) and diluted to an OD of 1.0 followed by 2-4 hours of incubation at 28°C with 200 rpm shaking to induce virulence. Infiltrations into *Nicotiana benthamiana* were done by mixing the prepared *A. tumefaciens* cultures in the indicated ratios in the methods sections below.

**Formaldehyde cross-linking**

*N. benthamiana* leaves infiltrated with AVRPM3$^{b2/c2}$-FLAG were allowed to incubate for 3 days followed by infiltration with a 1% formaldehyde solution and incubated at room temperature for 30 minutes. The leftover cross-linking reagent in the leaf was quenched by infiltrating a 125mM glycine solution. Infiltrated tissue was left for at least 30 minutes or until the tissue was dry again. Equal amounts of sample were harvested in liquid $N_2$ and immunoblotted.

**Protein extraction and immunoblot analysis**

Agrobacterium carrying the vectors with our constructs of interest or the p19 silencing suppressor were mixed in a plasmid:p19 ratio of 1:1 and infiltrated into *N. benthamiana*. 3

days post infiltration (dpi), 6 leaf discs (6mm diameter) were collected and ground in liquid nitrogen after which 150µl of 2x Laemmli buffer (62mM Tris-HCl pH6.8, 10% glycerol, 2% SDS, 0.02% bromophenol blue, 1% β-mercaptoethanol) was added and samples were incubated at 70ºC for 10 minutes followed by centrifugation at 22,000xg for 5 min. supernatant of the whole cell lysate was separated on 16% or 4-20% SDS-PAGE and transferred to a nitrocellulose membrane (GE Healthcare, Chicago, Illinois, USA). Membranes were probed with primary anti-FLAG (M2, F3165, Sigma-Aldrich) and secondary HRP conjugated anti-mouse (sc-516102, LabForce)

**Hypersensitive response measurements**

NLRs and AVRs were cloned into the 35S promoter driven binary vector pIPKB004 using gateway cloning (Bourras et al., 2019). Agrobacterium carrying the vectors with our constructs of interest or the p19 silencing-suppressor were mixed at a ratio of NLR:AVR:p19 of 1:4:1 and NLR:p19 of 1:1. Imaging of cell death induced by the hypersensitive response was assessed at 5 days post infiltration by fluorescence imaging using the Fusion FXImaging System (Vilber Lourmat, Eberhardzell, Germany) with previously established settings (Praz et al., 2017).

**Co-immunoprecipitation**

FLAG and HA tagged variants of AVRPM3$^{b2/c2}$, AVRPM3$^{a2/f2}$, SVRPM3$^{a1/f1}$, AVRPM17 and constructs were cloned into the 35S promoter driven binary vector pIPKB004 using gateway cloning (Bourras et al., 2019; Lindner et al., 2020). Agrobacterium carrying the vectors with our constructs of interest or the p19 silencing suppressor were mixed in a ratio of bait:prey:p19 of 1:1:1 prior to infiltrations. Tissue was collected at 2-3 dpi and ground in liquid nitrogen followed by protein extraction from 350mg of ground tissue in 1.5ml of lysis buffer (50mM Tris-Cl, pH8.0; 150mM NaCl; 5.0mM $MgCl_2$; 2.0mM $Na_2S_2O_5$; 10% Glycerol; 5.0mM

DTT; 1.0mM EDTA; 0.5% IGEPAL CA-630 (Sigma-Aldrich); 1mM PMSF; 1x Protease inhibitor cocktail EDTA free (Roche) for 30 minutes at 4°C and constant rotation. Lysate was clarified by centrifugation at 22,000xg for 5 min after which the supernatant was transferred and centrifuged again at 22,000xg for 30 min. 100µl of supernatant was taken as sample input and mixed with 30µl of 5x Laemmli buffer and heated at 70ºC for 10 minutes followed by a centrifugation at 22,000xg for 5 minutes. 1.0ml of supernatant was mixed with 10µl of prewashed anti-FLAG (A36797, Thermo Fisher) or PierceTM anti-HA (88837, Thermo Fisher scientific) magnetic beads and incubated for 2-3 hours at 4°C with constant rotation. Beads were washed 5 times with 500µl of lysis buffer followed by elution in 50µl of 2x Laemmli buffer at 70°C for 10 min at 1,000rpm. Proteins from input and eluates were separated on 16% or 4-20% SDS-PAGE and transferred onto nitrocellulose membranes (GE Healthcare, Chicago, Illinois, USA) followed by detection using HRP conjugated primary anti-HA (clone 3F10, 12013819001, Roche) and primary anti-FLAG (M2, F3165, Sigma-Aldrich) and secondary HRP conjugated anti-mouse (sc-516102, LabForce).

**Luciferase complementation assay**

For *in vivo* luciferase complementation assay, genes encoding AVRs without the signal peptide were cloned into pDEST-NLuc-gw, pDEST-CLuc-gw, pDEST-gw-NLuc or pDEST-gw-CLuc (Gehl et al., 2011) via gateway cloning. Prior to infiltrations, Agrobacterium carrying the vectors with our constructs of interest or the p19 silencing suppressorwere mixed at a ratio of NLuc:CLuc:p19 of 1:1:1. At 3 dpi infiltrated *N. benthamiana* tissue was collected using a 6mm leaf discs punch and placed in incubation buffer (10mM MES-KOH pH5.6; 10mM $MgCl_2$). After collection, an equal volume of incubation buffer supplemented with 2mM Luciferin (BioVision, Milpitas, California, US) was added to a final Luciferin concentration of 1mM. following a 5 minute dark incubation, luciferase luminescence signal was imaged for 25 min at 100ms intervals using the Spark imaging system (Tecan, Männerdorf, Switzerland).

**Bi-molecular fluorescence complementation assay**

For *in vivo* bi-molecular fluorescence complementation assays, the AVRPM3$^{b2/c2}$ coding gene was cloned into pGTQL1221-yN and pGTQL1221-yC (Lu et al., 2010) via gateway cloning. free mRFP which was used as a cytoplasmic and nuclear marker was expressed by transforming A. tumefaciens with a destination gateway that adds an N-terminal mRFP fusion (pGWB455 from Nakagawa et al., 2007) without an insert, this allows for expression of “free” mRFP with a short C-terminal amino acid tail (mRFP-ITSLYKKAERET-STOP). Prior to infiltration, Agrobacterium carrying the vectors with our constructs of interest or the p19 silencing suppressor were mixed at a ratio of yN:yC:mRFP:p19 of 1:1:1:1. Confocal images of infiltrated *N. benthamiana* tissue were taken at 3 dpi using a Leica SP5 confocal laser scanning microscopy system (Leica, Wetzlar, Germany) with an Argon (456nm) and DPSS561 (561nm) laser. Emission of YFP was collected at 465-505nm and emission of mRFP was collected at 570-620nm.

**Structural modeling of effectors**

The amino acid sequences from the effectors AVRPM3$^{a2/f2}$, AVRPM3$^{b2/c2}$ and SVRPM3$^{a1/f1}$ without the signal peptide were subjected to structural modeling using Alphafold2 or Alphafold3 with standard settings (Jumper et al., 2021).

Supplementary table 1 **Alphafold3 models of AVRPM3 effectors and SVRPM3$^{a1/f1}$ monomers and dimers**

Alphafold3 predictions of AVRPM3$^{a2/f2}$, AVRPM3$^{b2/c2}$ and SVRPM3$^{a1/f1}$ as well as homo- and heterodimer prediction of each combination. The average pLDDT (pLDDT$_{AVERAGE}$) values are calculated as the mean of the pLDDTs of all atoms in the models. Ranking_score, ptm and iptm are taken directly from the Alphafold3 output.

| **Alphafold3 models** | **pLDDT$_{AVERAGE}$** | **ranking_score** | **ptm** | **iptm** |
|---|---|---|---|---|
| avrpm3a_pu7_monomer | 88.64 | 0.82 | 0.82 | null |
| avrpm3a_pu7_dimer | 79.4 | 0.2 | 0.49 | 0.12 |
| avrpm3b_a_monomer | 48.88 | 0.52 | 0.35 | null |
| avrpm3b_a_dimer | 48.52 | 0.48 | 0.47 | 0.45 |
| svrpm3a_94202_monomer | 85.9 | 0.82 | 0.82 | null |
| svrpm3a_94202_dimer | 87.09 | 0.74 | 0.81 | 0.72 |
| avrpm3b_a_svrpm3a_94202_dimer | 58.24 | 0.18 | 0.5 | 0.07 |
| avrpm3a_pu7_svrpm3a_94202_dimer | 77.71 | 0.2 | 0.53 | 0.12 |
| avrpm3b_a_avrpm3a_pu7_dimer | 60.0 | 0.52 | 0.52 | 0.41 |

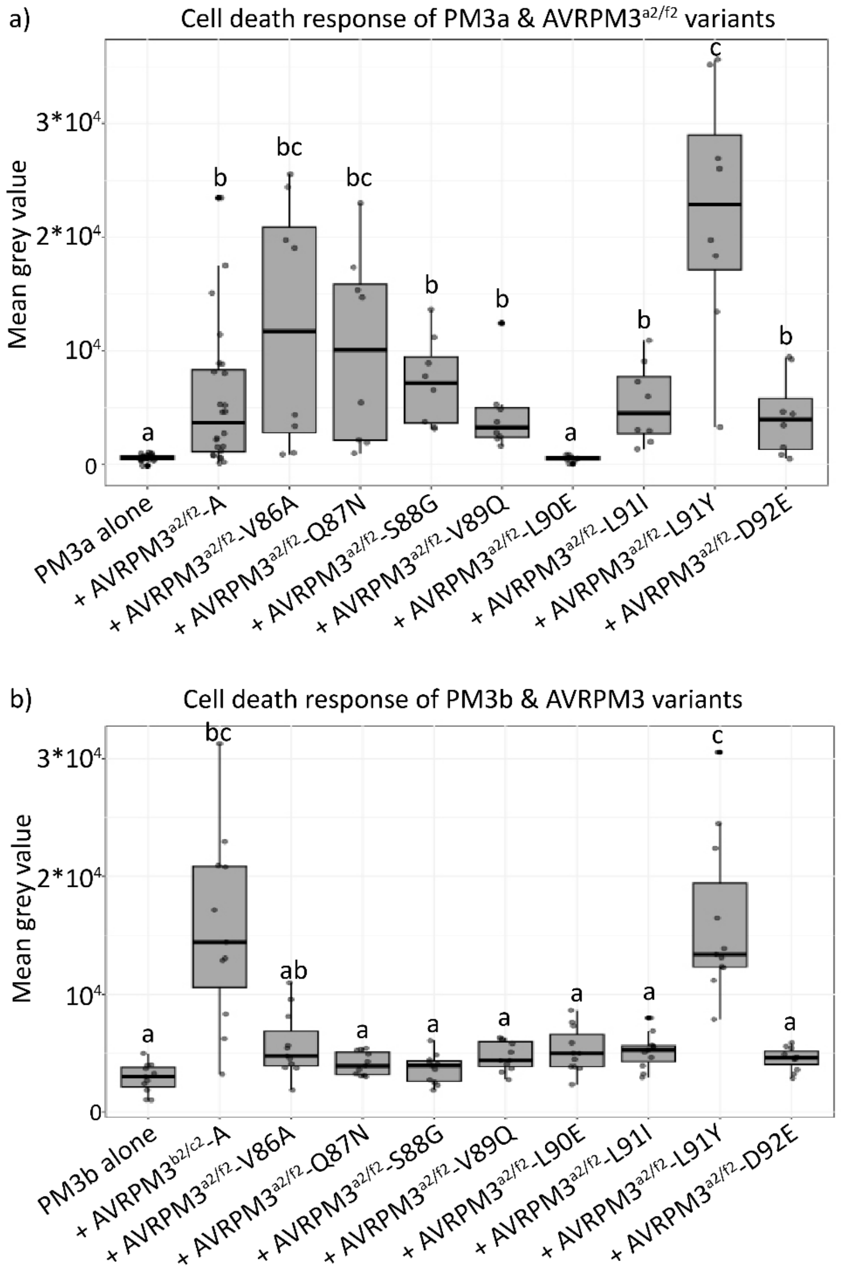


Supplementary figure 1 **A synthetic variant of AVRPM3$^{a2/f2}$ is recognized by the non-corresponding PM3b**

Cell death responses after transient Agrobacterium mediated transformation of *N. benthamiana* leaves. Fluorescence signal due to cell death was measured 5 days post infiltration. (a) PM3a was co-expressed with AVRPM3$^{a2/f2}$-A single amino acid substitution mutants lacking their signal peptides as indicated. PM3a was expressed alone or co-expressed with AVRPM3$^{a2/f2}$ lacking its signal peptide as negative and positive controls respectively. Fluorescence signal due to cell death was measured 5 days post infiltration. n=8 biological replicates per AVRPM3$^{a2/f2}$ mutant or n=24 biological replicates for AVRPM3$^{a2/f2}$-A and PM3a alone. (b) PM3b was co-expressed with AVRPM3$^{a2/f2}$-A single amino acid substitution mutants lacking their signal peptides as indicated. PM3b was expressed alone or co-expressed with AVRPM3$^{b2/c2}$-A lacking its signal peptide as negative and positive controls respectively. n=11 biological replicates for each sample. Statistical differences between the test and control were assessed using a pairwise Wilcoxon rank-sum test and a Benjamin-Hochberg p-value adjustment.

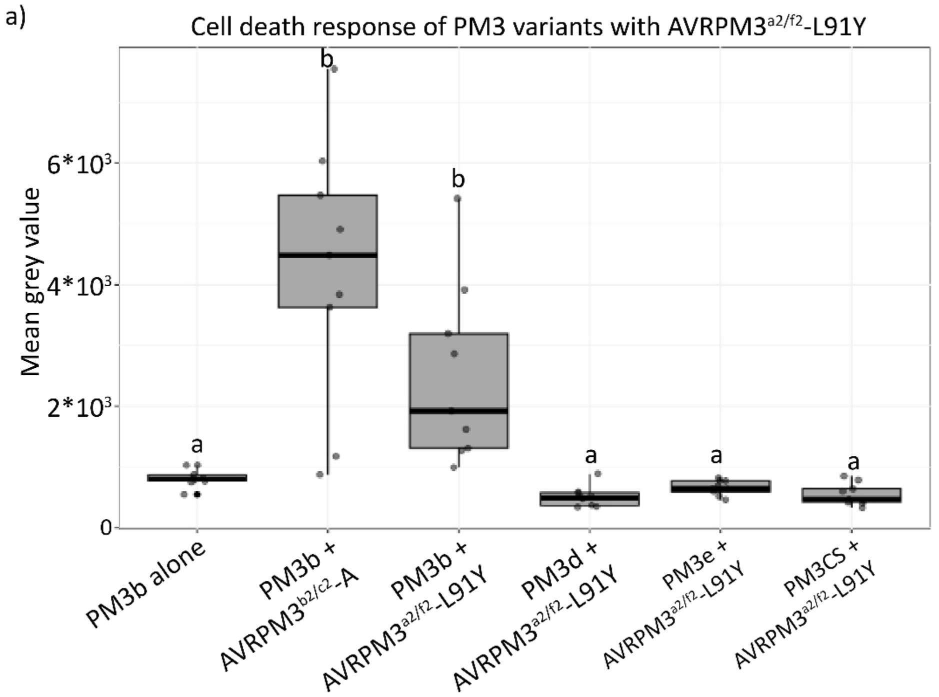


Supplementary figure 2 **AVRPM3$^{a2/f2}$-L91Y induced cell-death is specific to PM3b**

Quantification of the cell-death response at 5 days post infiltration in *N. benthamiana* induced by the co-expression of PM3 variants with AVRPM3$^{a2/f2}$-L91Y. PM3b co-expressed with variant A of AVRPM3$^{b2/c2}$ was used as a positive control. PM3b alone and PM3cs (a susceptible variant) co-expressed with AVRPM3$^{a2/f2}$-L91Y were used as negative controls. Fluorescence signal due to cell death was measured 5 days post infiltration with n=9 biological replicates. P-values were calculated using an un-paired Wilcoxon rank sum test.
